# CandyCollect: An Open-Microfluidic Device for the Direct Capture and Analysis of Salivary-Extracellular Vesicles

**DOI:** 10.1101/2024.10.09.617508

**Authors:** Corinne A. Pierce, Natalie P. Turner, Ingrid H. Robertson, Dakota McMillan, Xiaojing Su, Daniel B. Hatchett, Albert Shin, Olufemi Lijadu, Damon Wing Hey Chan, Sarah Feng, Karen N. Adams, Kezia Suryoraharjo, Alana Ogata, Erwin Berthier, Sanitta Thongpang, John R. Yates, Ashleigh B. Theberge, Lydia L. Sohn

## Abstract

Extracellular vesicles (EVs) are promising biomarkers for disease detection using a ‘liquid biopsy’ approach, in which they are enriched and analyzed directly from biofluids. However, implementing EV biomarker technologies in the clinic remains limited by the need for practical and patient-centric biofluid collection methods that are compatible with downstream EV processing and analysis. While saliva offers a non-invasive source of EVs, its complexity and heterogeneity—cells, debris, and other non-EV proteins—can present hurdles when using traditional analytical platforms. Here, we present the CandyCollect, a lollipop-inspired sampling device with open microfluidic channels, as a patient-friendly approach for rapid salivary EV capture. CandyCollect simplifies sample preparation by effectively pre-concentrating EVs in oxygen-plasma treated open microfluidic channels. In this proof-of-principle study, we show that following a 3-5 minute-oral sampling period, EVs collected by the CandyCollect can be released with high purity within minutes and subsequently quantified and analyzed for cargo content. We observed consistent EV capture across repeated collections within individuals and expected variability across healthy participants. Additionally, single and pooled collections of EVs from a healthy participant resulted in a concordant protein profile. Overall, the CandyCollect is a new platform for rapid, non-invasive salivary EV collection and analysis for clinical diagnostics.

## 1. Introduction

Extracellular vesicles (EVs) are lipid-bilayer enclosed nanoparticles continuously shed by cells (*1-3*) into the extracellular space. EVs contain a molecular suite of DNA, mRNA, miRNA, other RNAs, and proteins in their cargo (*4, 5*) that reflects a distinct signature linked to their cell of origin (*6-8*). EVs facilitate intercellular communication (*9*), intracellular waste removal (*10*), and a range of biological processes, from development and homeostasis (*11*) to cancer evolution and metastasis (*9*). While much research has focused on understanding the biogenesis and function of EVs (*12*), recent attention has pivoted to harnessing EVs’ clinical potential as biomarkers for a range of diseases (*13-17*), including cancer and neurodegenerative disorders (*18-20*).

Blood, urine, saliva, and cerebrospinal fluid (CSF) are replete with EVs at concentrations ranging from 10^6^ EVs/mL in CSF to 10^7^-10^9^ EVs/mL in saliva or urine to 10^10^ EVs/mL in blood (*21-26*), and EV biomarker discovery studies utilizing these sample types have been explored extensively (*17, 27, 28*). Despite great promise, a significant hurdle to employing EVs clinically is their efficient isolation for subsequent analysis, which typically requires EV-enriched samples of high purity and sufficient yields for downstream assays, both of which come with their own unique challenges (*29*). Traditional EV isolation methods include size and/or density-based strategies such as ultracentrifugation (UC) (*30, 31*), density gradient ultracentrifugation (DUC) (*32, 33*), size-exclusion chromatography (*34*), and affinity or charge-based separation (*35*). Although extensively used for EV-enrichment, these methods are not without challenges, including high-volume sample input requirements, low EV yields and compromised integrity, expensive instrumentation, labor intensive workflows (*36-39*), and lengthy processing times, particularly with UC and DUC (*33*). More recent methods of EV isolation (*40-46*) have addressed some of these challenges. For example, microfluidic platforms that utilize viscoelastic fluidics (*47*) or acoustofluidics (*46*) have demonstrated high EV yields (87-99%), low cost, ease of use, and preservation of EV integrity (*48*). However, purity and recovery, sample input, throughput, and incompatibility with some subsequent analysis methods still pose ongoing issues for these and many other contemporary methods (*34, 37-39, 49-56*). Similarly, challenges exist in the analytical tools used for EV biophysical characterization. Because of EVs’ nanoscale size, these tools are in the high-sensitivity realm and include transmission electron microscopy (TEM) (*57*), nanoparticle tracking analysis (NTA) (*58-60*), dynamic light scattering (*61*), and flow cytometry (*62, 63*), to name only a few. Additionally, they are constrained by high equipment costs and low throughput analysis. Overall, cost-effective and more efficient platforms that enable high recovery and purity of isolated EVs that do not impede subsequent analysis are crucial if EVs are to achieve their full potential as clinical biomarkers for disease.

Just as there is a challenge in developing an EV technology for efficient capture and analysis, so too is there a challenge in choosing the appropriate biofluid from which to isolate EVs. Among the aforementioned biofluids, saliva holds significant promise. Compared to blood and CSF, saliva can be easily collected non-invasively in a clinic or physician’s office or remotely at home. Yet, saliva is a complex, heterogenous biofluid consisting of cells, debris, proteins, enzymes, and mucins (*64, 65*) that make EV isolation challenging. Here, we employ the CandyCollect (*66-71*), a novel lollipop-inspired, saliva sampling device consisting of open microfluidic channels pretreated with oxygen plasma, to capture directly salivary EVs for enumeration and cargo analysis. Previously used to capture respiratory pathogens and commensal bacteria in saliva (*66-69*), we demonstrate that the CandyCollect effectively concentrates salivary EVs onto its surface within a 3-to 5-minute window. We further demonstrate that we can release the EVs from the CandyCollect with high purity also within minutes for either enumeration or cargo analysis. For enumeration, we employ a method we previously used to quantify virus and liposomes (*72*): specifically, we tag the EV lipid membranes with oligonucleotides (oligos) and subsequently perform qPCR to amplify and quantify the oligo tags. For cargo analysis, we employ mass spectrometry (MS)-based proteomics for the detection of canonical EV protein markers and assessment of inter-sample variability of EV proteins across repeated collections from the same individual.

## 2. Results

### 2.1. CandyCollect design and working principle of EV capture

The CandyCollect device is a lollipop-shaped polystyrene stick that has open microfluidic channels milled or injection molded into the “lollipop” end (**Figure 1** and **Materials and Methods**). The lollipop end is treated with oxygen plasma (*73, 74*), which has been shown to facilitate adhesion to salivary bacteria in human subject studies (*67, 68*). While the back and sides of the lollipop are coated with isomalt candy, the face of the device is left bare, leaving the plasma-treated microfluidic spiral open for saliva to flood the channel and for the subsequent continuous capture and accumulation of EVs (**Figure 1A**). The thickness of the isomalt candy on the CandyCollect device serves as a built-in incentive for sampling (*66*), encouraging participants to incubate the device in their mouth for a set period of time.

**Figure 1.**
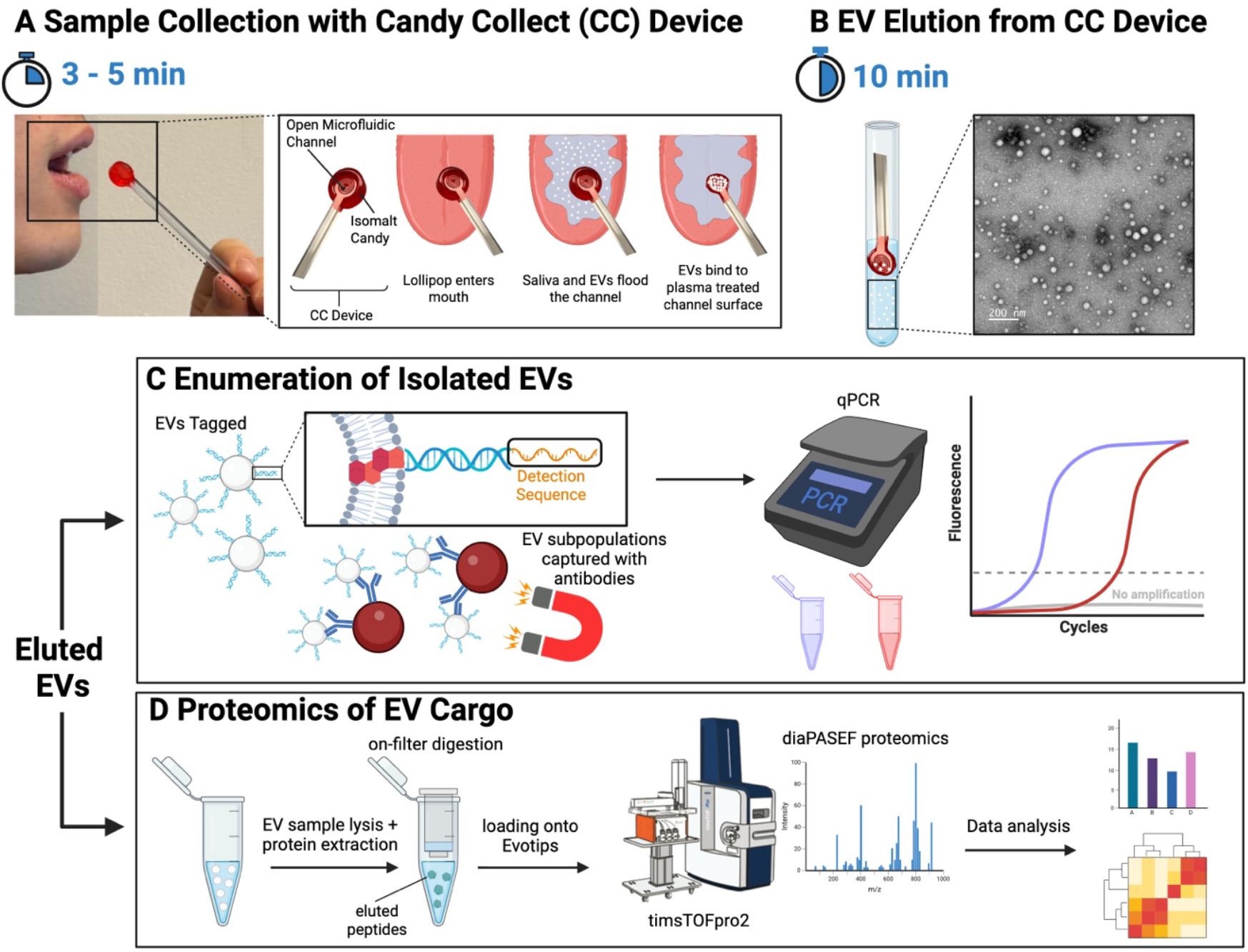
Salivary EV isolation and analysis workflow. A) CandyCollect device with a spiral open fluidic channel for EV capture and time-release isomalt/strawberry flavor (red). B) EVs are released from CandyCollect using Qiagen exo-Easy Maxi Kit elution buffer. Inset: TEM image of eluted EVs. Size bar = 200 nm. C) EVs are captured with antibody-coated paramagnetic beads and excess oligos are washed away. qPCR in which the detection oligo is amplified enabling measurement of the relative number of EVs. D) Mass spectrometry-based protein analysis digests proteins released from the EVs with trypsin followed by DIA-PASEF on a Bruker TIMS-TOF HT tandem mass spectrometer. Raw tandem MS data is processed using DIA-NN to obtain peptide and protein quantities at 1% FDR, with further processing for quantitative and qualitative analysis performed in RStudio.

Following incubation in the mouth, the CandyCollect device is immediately submerged into a Qiagen exoEasy Maxi Kit elution buffer for 10 min to release the EVs bound to the open microfluidic channels of the CandyCollect device (**Figure 1B**). After performing a buffer exchange into phosphate buffered saline (PBS) solution (**Materials and Methods**), the eluted EVs are ready for enumeration (**Figure 1C**) or cargo analysis (**Figure 1D**).

### 2.2. Characterization of Captured/Eluted EVs

We have examined the size and morphology of the captured/eluted EVs via TEM. Particle size analysis of three independent grid areas using the semi-automated image analysis platform TEM ExosomeAnalyzer (*75*) showed a consistent population of particles within the 30 - 60 nm size range, with most particles measuring 35 - 40 nm (**Figure 2A** and **Materials and Methods**). Although standard TEM is known to underestimate the size of vesicles due to the ‘shrinkage effect’ from sample fixatives that dehydrate vesicular structures (*76, 77*), these results confirm that the CandyCollect device isolates vesicles that fall within the size range of small EVs (30 - 150 nm). NTA of captured/eluted EVs (**Materials and Methods** and **Table S1**) determined an average concentration of 8.7 × 10^9^ EVs/mL per CandyCollect.

**Figure 2.**
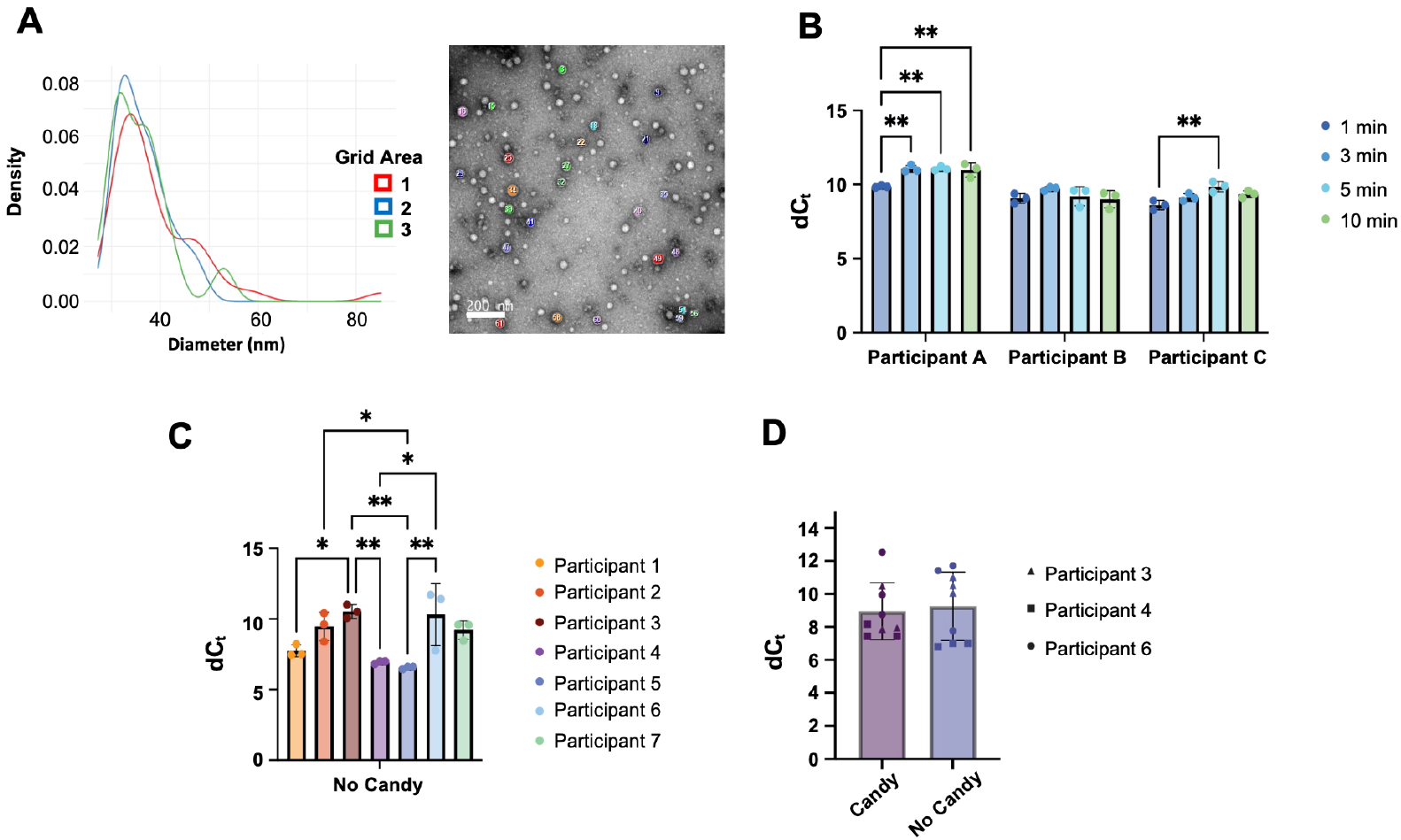
EV characterization, incubation time optimization, and quantification of EVs from healthy participants. A) Particle size distribution analysis from TEM imaging. Left panel shows kernel density plots of particle diameter measurements from three different grid areas (Grid area 1, n=42; Grid area 2, n=25; Grid area 3, n=30). Particle diameters were measured using TEM ExosomeAnalyzer following image preprocessing in ImageJ. Right panel shows a representative TEM micrograph with numbered particles identified for size analysis. Scale bar = 200 nm. B) The relative number of salivary EVs isolated from three healthy human participants using the CandyCollect platform. Each participant sucked on a total of twelve CandyCollects without isomalt candy: three for 1 minute (dark blue), three for 3 minutes (medium blue), three for 5 minutes (light blue), and three for 10 minutes (green). EVs were then eluted off of each CandyCollect and enumerated using the oligo labeling and subsequent qPCR-based method described in **Fig. 1**. There was no significant difference between 3 and 5 min CandyCollect incubation in a given individual. C) After a 5 min incubation in the mouth with 3 CandyCollects without candy, EVs were eluted off of each CandyCollect device and enumerated using the oligo labeling and subsequent qPCR-based method. As expected, there is significant differences in the relative number of EVs among participants, but little variation from device to device in a single participant. D) Comparison between candy-coated and bare CandyCollect devices. Participants sucked on three isomalt candy-coated and three candy-free CandyCollect devices, each for 5 min. EVs were eluted off of each CandyCollect device and enumerated using our oligo labeling and subsequent qPCR-based method. No significant differences in the number of EVs were observed with or without isomalt candy. For panels (B-D), the qPCR signal is presented as dC_t_, the difference in threshold cycle between each sample and a PBS control. Error bars = standard deviation. *: p<0.05; **:p<0.01, Two-Way ANOVA followed by Tukey multiple comparisons tests where the time factor accounted for 65% of the variance and the participants accounted for 16% (B), One-Way ANOVA followed by Tukey multiple comparisons (C), ns = no significance (p=0.72), two-tailed paired t-test (D).

### 2.3. Enumeration of isolated EVs using oligo-tagging and qPCR

Isolated EVs are labeled with oligos using our previously published method (*72*). Specifically, a 3’ cholesterol-tagged single-stranded DNA oligo (universal anchor) that had been prehybridized to a qPCR detection oligo self-embeds into the EV lipid bilayer membrane. A 5’ cholesterol-tagged universal co-anchor oligo is then introduced, which subsequently self-embeds into the EV lipid bilayer membrane and hybridizes with the universal anchor, thereby stabilizing the complex (**Figure 1C** and **Materials and Methods**). Labeled EVs are next isolated with paramagnetic beads (Dynabeads) functionalized with anti-CD63 antibody (Biolegend). CD63 was targeted, as it is a pan-EV marker (*49*). However, the generality of the method enables the use of beads functionalized with other antibodies. qPCR with primers specific for the detection oligos was performed on prepared labeled EV samples, thereby measuring the relative number of EVs (**Figure 1C** and **Fig. S1**). See **Materials and Methods** for details.

### 2.4. CandyCollect device effectively captures EVs after 3-5 min sampling time in the mouth

To determine the minimum sampling time needed to achieve saturation of EVs in the CandyCollect device, three healthy human participants were instructed to suck on a total of twelve candy-free CandyCollect devices each: three at 1 min, three at 3 min, three at 5 min, and three at 10 min (**Figure 2B** and **Materials and Methods**). To mitigate the introduction of any extra variables, sample collection for each experiment was conducted in the morning, between the hours of 9 am and 11 am, and at least 30 min after eating or drinking. No breaks were taken between CandyCollects used. To minimize bias, each participant was asked to incubate the CandyCollect devices in their mouth in different orders of sampling time: Participant C completed sample collection in descending order of incubation time (starting with 10 min and finishing with 1 min), Participant B completed sample collection in ascending order of incubation time (starting with 1 min and finishing with 10 min), and Participant A completed sample collection in a random order (i.e., starting with 5 min, then 3 min, then 10 min, and finally 1 min). EVs were immediately eluted off of the CandyCollect and analyzed using the qPCR method described above and in **Figure 1C**. To ensure we quantified only oligo labels associated with EVs, we considered dC_T_ between each sample and a PBS control that was processed in parallel and used as a background correction.

Across all three participants, a 1-min CandyCollect incubation led to significantly fewer EVs being captured as compared to the longer incubation times. CandyCollects that were incubated in the mouth for 3 min, 5 min, and 10 min, however, all captured nearly equivalent numbers of EVs, suggesting that a 3-5 min incubation is sufficient for EV saturation. Consequently, for all experiments subsequently performed, CandyCollects were incubated for 5 min in the mouth.

### 2.5. EV capture is robust among individuals

To investigate how robust EV capture with the CandyCollect is across individuals, seven healthy human participants were instructed to suck on three candy-free CandyCollect devices for 5 min each. EVs were immediately eluted off of each CandyCollect and enumerated as described in **Figure 1C**. Each data point (colored dot) at the top of each bar in **Figure 2C** is representative of one individual CandyCollect device from a given participant. As expected, there is a significant difference in EV capture across participants. Importantly, however, there is less variation in EV capture between devices within a given participant, though some within-participant variability is observed as expected in human subjects studies.

### 2.6. Isomalt candy does not interfere with EV capture or enumeration

To determine whether isomalt candy interferes with EV capture and subsequent enumeration, three participants were instructed to suck on six CandyCollects each: three that were candy-free and three that were coated with 0.5 g of strawberry-flavored isomalt candy (**Materials and Methods**). For the candy-coated CandyCollects, participants were instructed to remove the device from the mouth after exactly 5 min, regardless of whether all the candy had completely dissolved. Whereas for most participants, this amount of time was sufficient to dissolve the 0.5 g of isomalt candy, some participants could not dissolve all of the candy in the allotted time. All devices including those with trace amounts of isomalt candy were processed identically according to our method as described in **Figure 1C**. As **Figures 2D** and **S2** show, there is no significant difference in EV capture by devices with or without isomalt candy (p = 0.72). Thus, the presence of candy does not inhibit or influence EV enumeration using this qPCR-based method when using the CandyCollect.

### 2.7. Proteomic analysis of captured EV cargo reveals enrichment of canonical EV protein markers

EVs from a healthy 29-year-old participant were collected using, and subsequently eluted from, single and multiple pooled CandyCollects, and subjected to liquid chromatography tandem mass spectrometry (LC-MS/MS) **(Figures 1D** and **3**). Two-CandyCollect (2CC) pooled samples yielded more peptide precursors than One-CandyCollect (1CC) samples at 1% false discovery rate (FDR) (∼16,000 vs ∼12,500 in 2CC and 1CC, respectively), however both 1CC and 2CC resulted in identification of > 2000 protein groups, with gene set enrichment analysis showing significant enrichment of the terms ‘extracellular vesicle’, ‘extracellular exosome’, and ‘membrane-bounded vesicle’ (*p* < 1e-70). Multiple canonical EV markers were detected, including tetraspanins (CD81, TSPAN6), ESCRT-associated proteins (ALIX/PDC6I, CHMP4B, VPS35), flotillin-2, syntenin-1 (SDCB1), neutrophil-specific EV marker S100A8/A9, and an extensive catalog of RAB GTPases and SNARE proteins involved in vesicle trafficking (*78-80*). The presence of multiple heat shock proteins (HSPA8, HSP90, HSPA5) and annexins (ANXA1-6, ANXA11) further supports enrichment of multiple classes of EVs based on current community guidelines for biomolecular characterization (*81*). Salivary proteins were also detected, including mucins and amylases, alongside substantial neutrophil-derived content (MPO, PRTN3, defensins) and immunoglobulins. Collectively, these findings confirm effective EV enrichment from saliva, capturing a heterogeneous population likely comprising exosomes, microvesicles, and possibly apoptotic bodies, with cargo reflecting both the cellular origins and functional roles of salivary EVs in mucosal immunity and tissue homeostasis. Future work will include more nuanced differentiation of EV subtypes as well as investigation of non-EV associated proteins (e.g., mucins and amylases) that may provide complementary biological insight.

**Figure 3.**
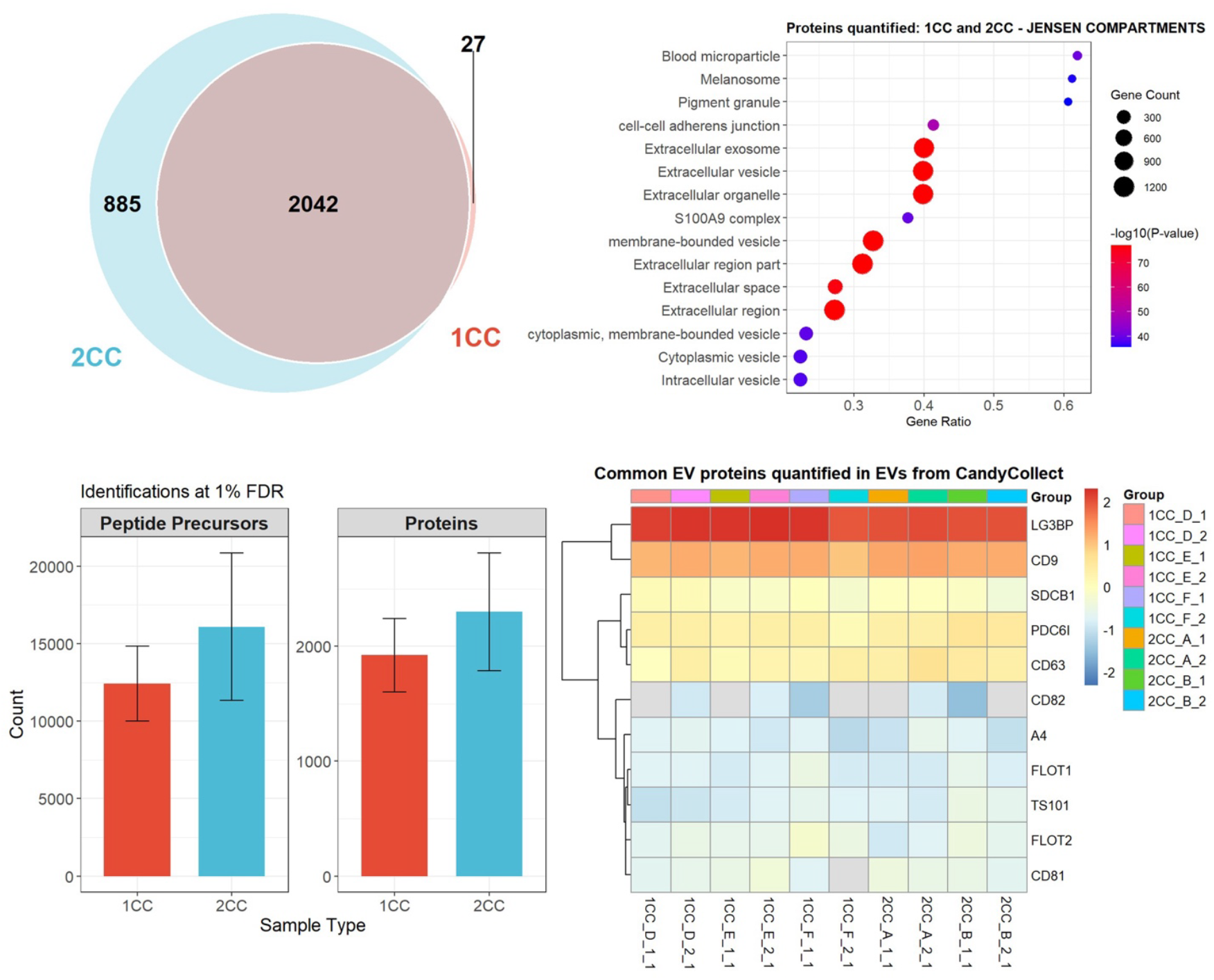
Proteomics analysis of CandyCollect (CC) sample replicates. Top left: Venn diagram of proteins identified in samples: 1CC (27), 2CC (885), and overlap (2042), showing that that a single CC is sufficient for most proteins and captures multiple EV protein markers (see heatmap), and that two samples can be combined if desired for less-abundant proteins but may also increase the detection of non-EV/contaminant proteins. Bottom left: Mean +/-SD of proteins and peptide precursors identified by DIA-NN at 1% FDR in 1CC and 2CC samples. Top right: Gene set enrichment analysis results from EnrichR against the Jensen COMPARTMENTS database for overlapping proteins in 1CC and 2CC samples (Venn: 2042). The top 15 enriched terms include ‘extracellular vesicle’, ‘membrane-bounded vesicle’, and ‘extracellular exosome’ (*p* < 1e-70). Bottom right: Heatmap of established EV protein markers quantified in CC samples (membrane proteins: CD9, CD81, CD82, CD63, FLOT1/2. Intraluminal proteins: TS101, PDC6I, SDCB1, LG3BP) and amyloid beta precursor protein (A4). Data were scaled by column to visualize intrasample variability and stability of EV protein signatures in CC samples.

The 885 proteins uniquely detected in the pooled (2CC) EV preparation likely represent lower-abundance EV cargo that falls below the detection threshold with a single collection (*82*). This includes additional ESCRT and endosomal sorting machinery (CHMP1B, CHMP3, CHMP4A/C, VPS28, VTA1, HGS) suggesting that pooling provides sufficient material to detect the vesicle biogenesis machinery itself, which may be present at lower copy numbers than abundant cargo proteins (*83*). Similarly, the unique detection of early endosomal regulators (EEA1, APPL1/2) (*84*), and autophagy markers (LC3A/B, GABARAPL2) (*10*) points to improved sensitivity for regulatory and trafficking components. The expanded mitochondrial (NDUF subunits, TCA cycle enzymes) and ER signatures (Sec proteins, TMED family, EMC components) may reflect either genuine detection of organelle-derived vesicles or simply that increased input unmasks these lower-abundance contaminants. The presence of additional signaling receptors (EGFR, ERBB3, EPH receptors) (*85*) suggests pooling enables detection of functionally relevant but scarcer EV subpopulations. Further analysis of sample collection variability was performed by examining the true detection rates (i.e., without filtering of missing values) and the relative abundance of canonical EV markers within samples (intrasample variation; **Figure 3**: Heatmap). Both 1CC and 2CC samples displayed consistent identification and stable relative quantities of EV proteins LG3BP, CD9, SDCB1, PDC6I, CD63, FLOT1, TS101, FLOT2. Amyloid precursor protein (A4) was assessed alongside canonical EV markers for its known association with Alzheimer’s Disease and displayed a similar inter-sample quantification pattern across 1CC and 2CC collections, suggesting that a single CandyCollect can capture EVs of sufficient quantity and purity for robust detection of these proteins. The tetraspanins CD81 (9/10 samples) and CD82 (5/10 samples) were not as consistently detected across samples, which could suggest a decreased prevalence of CD81/CD82-positive EV populations obtained using the CandyCollect. Future studies utilizing samples collected from more individuals are needed to confirm whether this is a feature associated with the CandyCollect, or whether this may be due to inter-individual variability in salivary EV protein cargoes.

## 3. Discussion

We have demonstrated a novel saliva sampling device—the CandyCollect—that can capture EVs for enumeration and cargo analysis. Specifically, we show that the CandyCollect can effectively capture and concentrate EVs in a simple 5-minute incubation period in the mouth. Further, EVs can be eluted off of the device and labeled with cholesterol-tagged oligos whose sequence can be amplified by qPCR for EV enumeration. Finally, as proof-of-principle, we show that EV cargo analysis can be performed using MS-based proteomics to identify thousands of proteins with significant enrichment of canonical EV protein markers.

This foundational work presents a rich array of topics for future work, including characterizing the proteomic landscape in healthy individuals and across the life span and using the CandyCollect device for population-level and multi-time point screening from populations that may be at risk for certain health conditions such as dementia or other chronic conditions that often present with molecular signatures that precede the symptomatic phase. Salivary EVs hold great potential to serve as a “molecular window” into other organs and tissues in the body, and the CandyCollect patient-centric sampling experience and simplified preanalytical workflow may allow for more frequent monitoring and subsequent high-content multiomic analysis, enabling tailored prediction of disease trajectories and treatments. Overall, the combination of CandyCollect sampling and EV analysis has the potential to advance our understanding of complex biomolecular mechanisms biomedical and become a powerful tool for clinical research.

## 4. Materials and Methods

### 4.1. Fabrication and Preparation of CandyCollect Devices

#### 4.1.1. CandyCollect device milling

A DATRON neo computer numerical control (CNC) milling machine, was used to mill the CandyCollect devices from 2 mm (GoodFellow, Cat# 235-756-86) and 4 mm (Goodfellow, CAT# 725-602-13) thick polystyrene sheets. After milling, the CandyCollect devices were sanded smooth, then sonicated with isopropanol (IPA) (FisherScientific, A451-4) and 70% v/v ethanol (FisherScientific, Decon™ Labs, 07-678-004)(*1, 2*). The engineering drawings of the 2 mm and 4 mm CandyCollect designs that are provided in **Figures S3** and **S4** were previously published in Lee *et al*. (*66*) and Tu *et al*. (*67*), respectively.

#### 4.1.2. CandyCollect device surface activation

CandyCollect devices were surface activated by plasma treating the devices with oxygen using a Zepto LC PC Plasma Treater (Diener Electronic GmbH Plasma Treater, Ebhausen, Germany). The following protocol is consistent with previous publications) (*66, 67*). Briefly, atmospheric gas in the chamber was removed until 0.20 mbar of pressure was reached. The chamber was then filled with oxygen gas until a pressure of 0.25 mbar was achieved and subsequently exposed to 70 W of voltage for 5 min (*66, 67*).

#### 4.1.3. CandyCollect devices with candy

Isomalt candy was added only to 4 mm-thick polystyrene CandyCollect devices where indicated. Lab members were trained in food safety and wore gloves and a mask to prepare and apply isomalt candy to CandyCollect devices. The isomalt candy was prepared as described in previous published papers (*66-68*). Briefly, heated isomalt was gradually added to water. The final addition of isomalt included food color. As the heated isomalt cooled, strawberry candy flavoring was folded into the mixture. The strawberry isomalt was poured onto a marble slab to set. During this time, the plasma-treated CandyCollect devices were cleaned using hot water and dish soap. Portions of the strawberry flavored isomalt candy were remelted. A silicone mold was used to apply 0.5 g of candy to CandyCollect sticks. After setting, CandyCollect devices with candy were moved to individual mylar bags (Belle KR, Cat# B0BZDCC8F8). Devices were stored in food-safe containers with a desiccant (Dry and Dry, Cat# B00DYKTS9C).

#### 4.1.4. CandyCollect devices without candy

CandyCollect devices without candy were cleaned by washing with hot water and soap. After drying, they were stored in food-safe containers. Participants used both 2 mm- and 4 mm-thick polystyrene devices based on availability of materials, but this difference in device thickness does not affect the channel geometry or EV capture.

### 4.2. EV Capture with, and Elution from, CandyCollect Device

CandyCollect devices, with and without isomalt candy, were incubated in a healthy participant’s mouth with the open microfluidic channel facing down on the tongue for 1, 3, 5, or 10 min. Following incubation, each CandyCollect device was immediately submerged in 400 µL of Qiagen exoEasy Maxi Kit (Cat# 76064) elution buffer for 10 min to release EVs bound to the open microfluidic channels of the CandyCollect device. The CandyCollect device was then removed from the elution buffer, and the buffer, itself, was transferred to a 0.5mL 10 kDa Amicon Ultra Centrifugal Filter (Millipore, Cat# UFC501024) and centrifuged for 15 min at 14,000 rcf at 4°C. After disposing the flow through, 400 µL of sterile phosphate buffered saline (PBS) was added to the filtration unit in order to perform a buffer exchange, as the Qiagen elution buffer is not compatible with the downstream oligo labeling step (described below). Centrifugation was performed again at 14,000 rcf at 4°C for 15 min. The Amicon filter was next transferred to a new collection tube and inverted before centrifuging at 1,000 rcf for 2 min at 4°C. Approximately 62 µL of EVs in PBS was recovered from the filter.

### 4.3. Transmission Electron Microscopy and Particle Size Analysis

Formvar-coated 300 mesh copper grids (Ted Pella Inc, product # 01753-F) were glow discharged (Easiglow, EMS, Hatfield, PA, USA) prior to use to create a positive surface charge for uniform distribution of EVs. EVs, in suspension (isolated from CandyCollects as described above) were pipetted onto a grid and allowed to settle for 60 seconds. After the solution was wicked away, 5 µL of 1% uranyl acetate in double-distilled water was added to the grid, for a total of 60 seconds. For some grids, 1x PBS was pipetted onto the grid surface and wicked away prior to the addition of the uranyl acetate stain to wash the grid or the grid of excess EV material. Excess uranyl acetate was wicked away and the grids were air dried for 3-5 min. Samples were imaged using a Tecnai 12 120kV TEM (FEI, Hillsboro, OR, USA) and data was recorded using a Gatan Rio 16 CMOS with Gatan Microscopy Suite software (Gatan Inc., Pleasanton, CA, USA).

TEM images were preprocessed in ImageJ (version 1.54g, NIH) to set the correct pixel size and convert images to 16-bit format. Particle diameter measurements were obtained using TEM ExosomeAnalyzer, a semi-automated computational platform designed for EV characterization (*75*). Default settings were used and manual corrections were applied as necessary to ensure accurate particle detection and size estimation. Measurements were collected from three different grid areas (*n* = 42, 25, and 30 particles, respectively).

Size distributions were visualized using kernel density estimation plots generated in R (RStudio 2026.01.0+392 “Apple Blossom” Release) using the ggplot2 package. Kernel density estimation provides a smoothed representation of the diameter distribution, where the area under each curve is normalized to one, allowing for direct comparison of distribution shapes across grid areas with different particle counts.

### 4.4. NTA Analysis

Three CandyCollect samples were obtained from one individual and processed as described above, with the additional step of 0.22 µm filtration through a cellulose acetate membrane filter (polypropylene housing; Corning, Cat# CLS8161) prior to molecular weight filtration, to reduce background noise prior to analysis. Samples were diluted 1:20 in PBS and analyzed using a NanoSight NS300 running NTA software version 3.4 (build 3.4.4). Capture settings were as follows: camera type: sCMOS; laser type: Blue 405; camera level: 15; slider shutter: 1206; slider gain: 366; FPS: 25; syringe pump speed: 50. Five videos of 1498 frames each were recorded per sample at temperatures ranging from 19.0–22.7°C. Analysis settings were as follows: detect threshold: 5; blur size: auto; max jump distance: auto (ranging from 12.7–26.3 pix across samples). Viscosity was set to water and ranged from 0.936–1.023 cP across samples, consistent with the observed temperature variation. Sample 1 yielded particle concentrations below the instrument’s recommended detection range and was excluded from further analysis; downstream analyses therefore reflect *n* = 2 samples. See **Table S1**.

### 4.5. Preparation and Functionalization of Colloids

1 µL of Dynabeads M-270 Streptavidin beads (Cat# 65305) per sample was pipetted into a nuclease-free microcentrifuge tube and washed with 200 µL of 0.2% bovine serum albumin (BSA) before being placed on a magnetic rack. After a pellet formed on the wall of the tube (due to the presence of a magnetic field), the buffer was carefully pipetted out before repeating two more 0.2% BSA washes. After the third and final wash, the beads were resuspended at a 1:10 ratio (10x volume originally pipetted from the colloid stock bottle) in 0.2% BSA and biotinylated antibody (anti-CD63 antibody, Biolegend Cat #353017) was added at 10 µg per mL of diluted colloids and incubated on a hula rocker for 30 min. Following incubation, the colloids were placed back on the magnetic rack, and the remaining solution was removed. Three more 0.2% BSA washes were performed before the colloids were again resuspended, this time with 3% BSA, and incubated on a hula rocker for > 30 min.

### 4.6. Cholesterol-Oligo Labeling of EVs and Capture of EVs on Colloids

Universal and co-anchor strands (*72, 86*) (**Table S2**) consisting of 41 and 20 nucleotides long, respectively, and 5’ and 3’ cholesterol tags were purchased from IDT. The detection strand, also purchased from IDT, is based on the Arabidopsis thaliana FLG22-inducted receptor-like kinase 1 (FRK1) gene because of its lack of sequence homology with human, viral, and bacterial genes, preventing unintended amplification with human samples (*72*). The length of the detection strand—130 nt long—was chosen to maximize PCR efficiency while also maintaining labeling efficiency. See Carey *et al*. (*72*) for additional information regarding oligo sequence design. See **Table S2** for the oligo sequences used.

Cholesterol-universal anchor and detection oligo were prehybridized at a 1:1 ratio (oligos were resuspended in Molecular Biology-Grade H_2_O at 100 µM to a final concentration of 50 µM) and incubated at 37°C for 15 minutes. This stock solution was further diluted 1:100 in nuclease-free H_2_O to obtain a 500 nM working solution. 100 µM co-anchor was diluted 1:200 to obtain a 500 nM working solution.

10 µL of EVs in buffer out of the total 62 µL that was recovered from each Amicon filter (as described above) was pipetted to fresh microcentrifuge tubes. 1 µL of 500 nM anchor + detection oligo solution was added to each EV sample and incubated at room temperature for 15 min. Following incubation, 1 µL of 500 nm co-anchor oligo solution was added to each EV sample. 10 µL of the prepared diluted functionalized colloid solution described above were added to each EV sample (each sample had a final volume of 22 µL) and incubated on hula rocker for 60 min. Following incubation, each sample was placed on a magnetic rack, and the remaining solution was removed. Each pellet was resuspended in 200 µL of 0.2% BSA and transferred to a fresh microcentrifuge tube. 0.2% BSA wash was repeated four more times with each sample being transferred to a new microcentrifuge tube with every wash before the pellets were resuspended in 26 µL of nuclease-free water.

### 4.7. qPCR

500 nM primers were prepared by diluting each forward and reverse FRK1 primers 1:100 in nuclease-free water. Mastermix consisted of 10 µL of PowerUp SYBR Green Master Mix (Applied Biosystems Cat# A25776) and 2 µL of 500 nM forward/reverse primer mixture per reaction. To each tube containing 26 µL of sample, 39.4 µL of master mix was added and mixed thoroughly before pipetting 20 µL to each well on the qPCR plate. The CFX Duet Real-Time PCR System (Bio-Rad CFX Maestro 2.3, software version 5.3.022.1030) was run according to standard (non-fast) parameters as specified in the PowerUp SYBR Green Master Mix handbook.

### 4.8. Mass Spectrometry

#### 4.8.1. Sample Preparation and LC-MS/MS Analysis

CandyCollect samples were processed for proteomic analysis using a modified filter-aided sample preparation (FASP) protocol for EVs (*87, 88*). In brief, samples were solubilized in a lysis buffer containing 1% *w/v* sodium deoxycholate (SDC) and 1% *w/v* sodium dodecyl sulfate (SDS), 100 mM dithiothreitol (DTT), and 100 mM Tris-HCl pH 8.5. Samples were loaded onto a 30 K molecular weight cut-off centrifugal filter (Nanosep™ Omega membrane, Cytiva; catalog number OD030C34) and centrifuged at 14,000 x*g* for 15 min at 21 °C with the flow-through discarded. On-filter protein denaturation was performed with 200 µL of 25 mM DTT in 8 M urea for 1 h at RT, followed by another round of centrifugation, and the flow-through discarded. Filters were washed once with 8 M urea and samples were alkylated with 100 µL of 50 mM iodoacetamide in 8 M urea by incubating in the dark for 20 min at RT. Filters were centrifuged and washed once with 400 µL of 8 M urea and once with 400 µL of 100 mM ammonium bicarbonate, and the flow-through discarded after each centrifugation step. Samples were digested overnight with trypsin suspended in 100 mM ammonium bicarbonate (0.1 µg/µL; enzyme:substrate ratio 1:50). The following day, peptides were eluted into clean Eppendorf protein lobind 1.5 mL sample tubes via centrifugation. Samples underwent one additional elution with 40 µL of 100 mM ammonium bicarbonate into the same tube. Peptide concentration estimation was performed using a colorimetric peptide assay (Pierce, Thermo Scientific, Catalog number 23275) according to the manufacturer’s instructions. Samples were loaded onto evotips (Evosep) according to the manufacturer’s instructions.

Five-hundred ng of peptides per sample were injected into the mass spectrometer in technical duplicate via elution from the column tip into a captive spray source, and peptides were nanosprayed into the MS. Samples were analyzed by parallel accumulation–serial fragmentation combined with data-independent acquisition (DIA-PASEF) (*89*) on a timsTOF HT mass spectrometer (Bruker) with a 15spd method (∼88 min gradient) at a flowrate of 220 nL/min. Buffer A consisted of H_2_O/0.1% FA and Buffer B was Acetonitrile/0.1% FA (LC-MS grade, Fisher Scientific). Peptides were separated by reversed-phase HPLC on an in-house packed analytical capillary column with integrated in-house pulled tip, with dimensions 25 cm, 150 nm internal diameter, and BEH 1.7 µm C18 resin (Waters). Capillary voltage was set to 1500 V and dry gas flow and temperature 3.0 L/min and 180 °C, respectively. DIA-PASEF MS1 scan range was 100 – 1700 *m*/*z* and ion mobility (1/K0) = 0.6 – 1.6 V s cm^2^. Accumulation and ramp time were 100 ms, and cycle time was 1.16 s. The PASEF MS/MS window scheme was set to 32 wide *m*/*z* isolation windows from 400 – 1200 *m*/*z* of width 25 m/z with 1 *m*/*z* overlap. MS2 TOF resolution was 45,000 and singly charged precursor exclusion enabled.

#### 4.8.2. MS Data Processing and Analysis

Raw data (.d files) were imported into DIA-NN v1.8.1 (*90*) and searched at 1% FDR against an *in silico* human predicted library containing 20,405 sequences with common contaminants appended. Carbamidomethylation on C was set as a fixed modification, trypsin was set as the digestion enzyme, and up to 1 missed cleavage was allowed. All other settings were as default. Further data processing performed in RStudio (v2025.05.0+496, R v4.5.0). Following common contaminant removal, results were filtered at Q.Value, PG.Q.Value, and Global.Q.Value < 0.01. Protein quantities were obtained using the diann_maxlfq function from the ‘diann’ R package (v1.0.1). Protein lists were filtered for gene set enrichment analysis by retaining proteins with < 2 missing values within each sample type (1CC or 2CC). The heatmap in **Figure 3** was created using the R package ‘pheatmap’ (v1.0.13), with scaling = ‘column’. Enrichment analysis was performed using the ‘EnrichR’ package (v3.4) against the Jensen COMPARTMENTS database. The number of peptide precursors and protein groups identified at 1% FDR were extracted directly from the DIA-NN statistics output file and plotted in RStudio (mean +/-standard deviation).

### 4.9. Statistical Analysis

Statistical analysis was performed with GraphPad Prism 10.3.1 software. Data were presented as mean +/-standard deviation. Statistical tests used in this paper included one-way and two-way ANOVAs with Tukey’s multiple comparison test and a paired t-test specific statistical tests, and sample sizes can be found in the appropriate figure captions. p< 0.05 was considered statistically significant.

## Acknowledgements

We thank members of the Theberge and Sohn Laboratories for helpful discussions. We also thank Reena Zalpuri and the staff at the University of California Berkeley Electron Microscope Lab for advice and assistance in electron microscopy sample preparation and data collection, and Evan Carpenter and the Randy Scheckman Laboratory for assistance in performing NTA analysis. We would also like to thank all the participants who gave their time to this study.

## Author Contributions

C. A. P., A. B. T., and L. L. S. conceived of the project. C. A. P., N. P. T., and D. M. performed all experiments presented. C. A. P., I. H. R., and N. P. T. analyzed the results. C. A. P. and X. S. performed initial experiments and developed capture and analysis protocols; X. S., I. H. R., and D. B. H. advised on CandyCollect procedures. I. H. R, D. B. H., A. S., D. C., and S. F. fabricated the CandyCollect devices. I. H. R. and K. N. A. advised on human subjects procedures and study design. K. S. and A. O. performed initial protein analysis and EV size characterization experiments. A. B. T., E. B., and S. T. provided leadership for the development, production, and implementation of the CandyCollect device. O. L. designed Figure 1. C. A. P., N. P. T., J. R. Y., B. T., and L. L. S. wrote the manuscript with input and critical review from the other authors.

## Funding

National Institutes of Health grant R35GM128648 (A. B. T.)

Schmidt Sciences, LLC via a Schmidt Science Polymath Award (A. B. T.)

National Institutes of Health grant R01EB024989 (L. L. S.)

Agilent Biodesign Gift (L. L. S.)

National Institutes of Health grant 5R01MH100175-11 (J. R. Y.)

National Institutes of Health grant 2P60 AA006420-40 (J. R. Y.)

National Institutes of Health grant 5T32AA007456-42 (N. P. T.)

## Conflict of Interests

Ashleigh B. Theberge, Xiaojing Su, Erwin Berthier, and Sanitta Thongpang filed patent 63/152,103 (International Publication Number: WO 2022/178291 Al) through the University of Washington on the CandyCollect oral sampling device. Erwin Berthier and Ashleigh Theberge filed patent 63/683,571 through the University of Washington on a related platform. Ashleigh B. Theberge reports filing multiple patents through the University of Washington and receiving a gift to support research outside the submitted work from Ionis Pharmaceuticals. Erwin Berthier is an inventor on multiple patents filed by Tasso, Inc., the University of Washington, and the University of Wisconsin. Sanitta Thongpang has ownership in Salus Discovery, LLC, and Tasso, Inc. Erwin Berthier has ownership in Salus Discovery, LLC, and Tasso, Inc. and is employed by Tasso, Inc. However, this research is not related to these companies. Sanitta Thongpang, Erwin Berthier, and Ashleigh B. Theberge have ownership in Seabright, LLC, which will advance new tools for diagnostics and clinical research, including the CandyCollect device. The terms of this arrangement have been reviewed and approved by the University of Washington in accordance with its policies governing outside work and financial conflicts of interest in research. Lydia L. Sohn is an awardee of US Patent No. 12,229,291: “Lipid-DNA Labeling of Extracellular Vesicles for Amplification Quantitation,” with T. Carey and M. Kozminsky, issued June 24, 2025. The other authors have no conflicts of interest to disclose.

## Data availability

The data needed to evaluate the conclusions in the paper are present in the paper and/or the Supplementary Materials. The mass spectrometry proteomics data have been deposited to the ProteomeXchange Consortium (http://proteomecentral.proteomexchange.org) via the PRIDE partner repository (*91*) with the dataset identifier PXD078068.

## Ethics Statement

Human subject studies were conducted under an IRB-approved protocol (UC Berkeley Committee for Protection of Human Subjects, Protocol ID: 2024-07-17631), and all subjects provided written informed consent prior to taking part in the study.

## Notes

http://proteomecentral.proteomexchange.org

### Summary of Updates

Additional co-authors, a co-first author, and co-corresponding author. New title and abstract. Additional data (Fig. 2a; Fig. 3; and Supplementary Figures). Modified Figure 1.

