## Supplemental Information for "CandyCollect: An Open-Microfluidic Device for the Direct Capture and Analysis of Salivary-Extracellular Vesicles"

**Table S1. NTA measurement of EVs eluted from the CandyCollect.**

| | CC1 | CC2 | CC1 + CC2 avg |
|---|---|---|---|
| **Concentration (particles/mL)** | 1.07 x $10^{10}$ | 6.02 x $10^{9}$ | 8.37 x $10^{9}$ |

**Table S2. Oligonucleotide and primer sequences.[a]**

| Name | Modification | Sequence (5'-3') | Annotation |
|---|---|---|---|
| Universal Anchor | 3' Cholesterol-TEG | TGGAATTCTCGGGTGCCAAGGGAATTC GTAACGATCCAGCTGTCACT | Detection adhesion sequence Co-anchor adhesion sequence |
| Universal Co-Anchor | 5' Cholesterol-TEG | AGTGACAGCTGGATCGTTAC | Anchor adhesion sequence |
| Detection Oligo | N/A | GAATTCCCTTGGCACCCGAGAATTCCA TGAAGGAAGCGGTCAGATTTCAACAGT TGTCGCTGGATCCATCGGTTGTTCTTCT TGAAGTGATTACAGGCCAACCTGCTAT TCAGTCAGTCAGTCAGTCAGT | Anchor adhesion sequence Detection sequence |
| FRK1 Forward Primer | N/A | CGGTCAGATTTCAACAGTTGTC | |
| FRK1 Reverse Primer | N/A | AATAGCAGGTTGGCCTGTAATC | |

[a] Name indicates distinct sequences, and colors indicate complementary regions that hybridize in our assay.

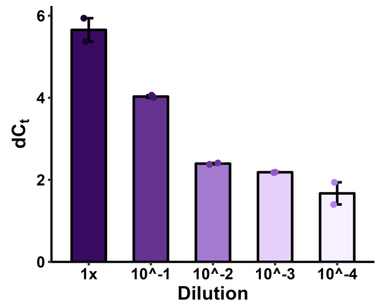


**Figure S1: Standard Curve for Oligo Tagging Method:** A five-point 1:10 serial dilution series was prepared from a CandyCollect EV sample, beginning at 1/6 of the total recovered volume (equivalent to the input volume used for a standard sample). Each dilution point was carried through the full cholesterol-oligo labeling and bead capture workflow independently, as described above, before qPCR analysis. Each dilution point was run in duplicate, and a no-template control (NTC) was included; NTC Cq values were sufficiently distinct from sample Cq values to confirm specificity.

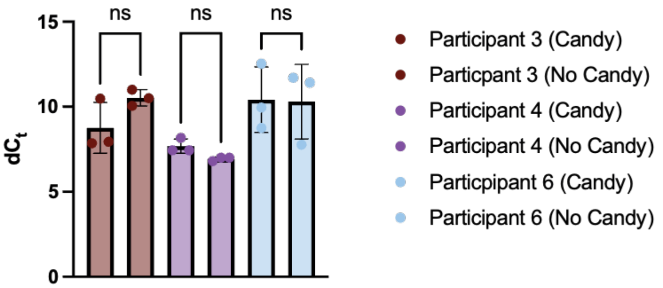


**Figure S2: Candy vs. No Candy CandyCollects**. Each participant sucked on a total of 6 CandyCollects, 3 with isomalt candy and 3 without, for 5 min each. EVs were eluted off of each CandyCollect and enumerated using our oligo-tagging method. The qPCR signal is presented as $dC_t$, the difference in threshold cycle between each sample and a PBS control. Error bars = standard deviation. A two-way ANOVA was used to analyze these data; between Candy vs No Candy there is no significant difference (p-value = 0.66), but significant differences were observed between participants as expected (p-value=0.0054) with an alpha of 0.05

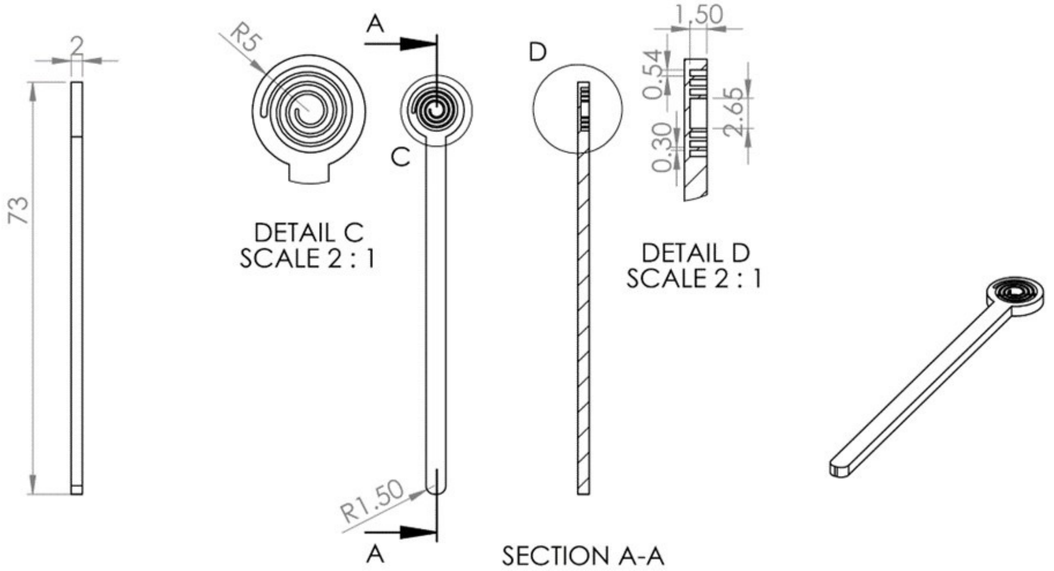


**Figure S3: CAD design of the CandyCollect platform**. Engineering drawing for the 2 mm CandyCollect device. Reproduced with permission from the authors (Lee *et al*. Figure S1).[1] All dimensions are in mm.

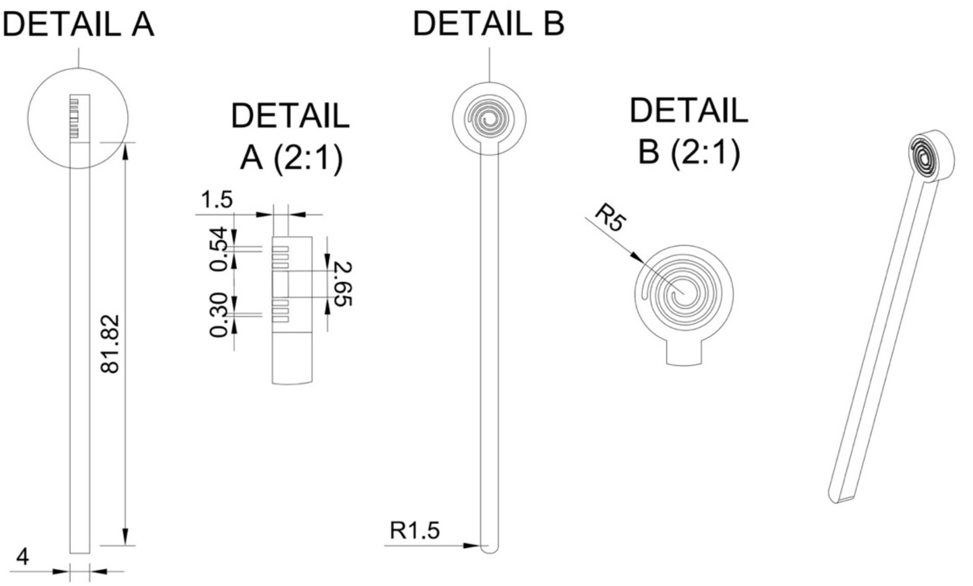


**Figure S4: CAD design of the CandyCollect platform.** Engineering drawing for 4 mm CandyCollect device. Reproduced with permission from the authors (Tu *et al*., Figure S1).[2] All dimensions are in mm.
